# Comparison of subtyping approaches and the underlying drivers of microbial signatures for chronic rhinosinusitis

**DOI:** 10.1101/494393

**Authors:** Kristi Biswas, Raewyn Cavubati, Shan Gunaratna, Michael Hoggard, Sharon Waldvogel-Thurlow, Jiwon Hong, Kevin Chang, Brett Wagner Mackenzie, Michael W. Taylor, Richard G. Douglas

## Abstract

Chronic rhinosinusitis (CRS) is a heterogeneous condition characterised by persistent sinus inflammation and microbial dysbiosis. This study aimed to identify clinically relevant sub-groups of CRS patients based on distinct microbial signatures, with a comparison to the commonly used phenotypic subgrouping approach. The underlying drivers of these distinct microbial clusters were also investigated, together with associations with epithelial barrier integrity.

Sinus biopsies were collected from CRS patients (n=23), and disease controls (n=8). Expression of 42 tight junction genes was evaluated using quantitative PCR, together with microbiota analysis and immunohistochemistry for measuring mucosal integrity and inflammation.

CRS patients clustered into two distinct microbial sub-groups using probabilistic modelling Dirichlet (DC) multinomial mixtures. DC1 exhibited significantly reduced bacterial diversity, increased dispersion, and was dominated by *Pseudomonas, Haemophilus*, and *Achromobacter*. DC2 had significantly elevated B-cells, incidence of nasal polyps, and higher numbers of *Anaerococcus, Megasphaera, Prevotella, Atopobium*, and *Propionibacterium*. In addition, each DC exhibited distinct tight junction gene and protein expression profiles compared with controls. Stratifying CRS patients based on clinical phenotypic subtypes (absence or presence of nasal polyps (CRSsNP or CRSwNP respectively) or with cystic fibrosis (CRSwCF)) did account for a larger proportion of the variation in the microbial dataset compared with DC groupings. However, no significant differences between CRSsNP and CRSwNP cohorts were observed for inflammatory markers, beta-dispersion and alpha diversity measures.

In conclusion, both approaches used for stratifying CRS patients had benefits and pitfalls, but DC clustering did provide greater resolution when studying tight junction impairment. Future studies in CRS should give careful consideration into the patient subtyping approach used.

**Importance:** Chronic rhinosinusitis (CRS) is a major human health problem that significantly reduces quality of life. While various microbes have been implicated, there is no clear understanding of the role they play in CRS pathogenesis. Another equally important observation made for CRS patients is that the epithelial barrier in the sino-nasal cavity is defective. Finding a robust approach to subtype CRS patients would be the first step towards unravelling the pathogenesis of this heterogeneous condition. Previous work has explored stratification based on clinical presentation of the disease (with or without polyps), inflammatory markers, pathology, or microbial composition. Comparing between the different stratification approaches used in these studies has not been possible due to different cohorts, analytical methods, or sample sites used. In this study, two approaches of subtyping CRS patients were compared and the underlying drivers of the heterogeneity in CRS were also explored.

## Introduction

As the first line of defence against inhaled antigens from the external environment, the upper airway epithelium represents an important physiological barrier (1). Maintaining the close cell to cell proximity that is central to the integrity of this barrier are tight junctions, which play a critical role in host defence (2, 3). Tight junctions comprise a range of transmembrane and scaffolding adaptor proteins that include occludin, claudins, junctional adhesion molecules, and zonula occludens (ZO) (4). These proteins limit the passage of macromolecules by sealing off the paracellular spaces between epithelial cells. On the other hand, in cases of tissue inflammation, the opening up of tight junctions assists in the release of tissue fluids and the influx of inflammatory cells and by doing so helps speed resolution. Accordingly, tight junctions are considered as the gatekeepers of inflammatory disease. Recent studies have considered the role of epithelial barrier defects in a number of chronic inflammatory conditions, including chronic rhinosinusitis (CRS) (5, 6). CRS is an inflammatory condition of the upper respiratory tract persisting for over 12 weeks, affecting 5% of the general population (7). Symptoms include nasal discharge, facial pain, loss of smell and headaches. CRS is a complex and heterogeneous disease, with many underlying factors that present with similar symptoms, making it challenging to separate CRS into clinically relevant subtypes. Traditionally, classification of CRS has been based on the clinical phenotypes – the presence (CRSwNP) or absence (CRSsNP) of polyposis. Approximately 25-30% of CRS patients present with nasal polyps (8), and this cohort is considered to have a Th2-predominated inflammatory response compared with idiopathic CRS without nasal polyps (CRSsNP) that has Th1-type responses. However, this simplified view misrepresents the true complexities of CRS.

Recent efforts have investigated the pathogenesis of CRS based on inflammatory markers (9, 10), microbiota composition (11, 12), pathology or clinical factors (13, 14), but these studies have produced little consensus on appropriate strategies to subtype patients. The role of microbes in CRS remains unclear due to the frequent lack of resolution with antimicrobial treatment. It has been suggested that a loss of overall microbiota diversity and deleterious community changes (collectively termed ‘dysbiosis’) are more characteristic of CRS patients than a single, disease-causing organism (15).

Stratification of patients based on probabilistic modelling of the bacterial communities in lower respiratory diseases such as asthma and HIV-infected pneumonia patients has been used successfully to classify immunological or clinical phenotypic variation across cohorts (16, 17). Using a similar approach, a recent study subtyped CRS patients based on their microbial community profiles (11). Each distinct microbial state was dominated by one bacterial family and associated with a unique clinical and host immune response. Previous studies have found that large interpersonal variations in the sino-nasal microbiome of CRS patients (subtyped based on clinical diagnosis) can make classification difficult (12). Accordingly, a distinct advantage of stratifying patients based on microbial community profiles is that it will help to resolve the microbial heterogeneity of CRS and is a step towards implementation of a precision medicine approach. The underlying drivers of these distinct microbial states in CRS are yet to be investigated.

Several studies have investigated epithelial integrity in CRS mucosa and found that CRS patients with nasal polyps (CRSwNP) have severely disrupted epithelia with decreased expression of occludin and ZO-1 (6, 18, 19). These efforts also suggest a role for cytokines (IFN-γ and IL-4) in disrupting tight junctions, whereas the influences of the microbes on the host are less understood.

One common hypothesis is that a loss of tight junction integrity in CRS patients could lead to the entry of environmental agents, including microbes, into host tissues (20). It remains unclear whether the microbes are the cause of tissue damage or if microbial patterns or associations are the consequence of a leaky epithelium, leading to a build-up of microbial cells in the tissue. The mechanism by which bacteria penetrate the epithelial barrier is unknown, but their presence in CRS tissue (in particular CRS with cystic fibrosis (CRSwCF)) presumably reflects a breakdown in mucosal integrity (21). Furthermore, any changes caused to the micro-environment (such as lack of mucociliary clearance, damaged surfaces for bacterial cell adherence) could have an impact on microbial composition and structure, as observed previously in gut studies (22).

In this study, we aimed to stratify CRS patients based on their microbial community composition using a probabilistic modelling approach and using the traditional phenotypic approach. In addition, we investigated several possible underlying influences on these microbial states by measuring gene and protein expressions of host tight junction, epithelial integrity and inflammatory cells (T-cells, B-cells, and macrophages) in the sino-nasal tissue biopsies.

## Results

### Clinical parameters

Non-parametric pairwise comparisons were made based on clinical factors in this cohort of 31 subjects (CRS = 23, disease controls = 8). Patient demographics are included in Table S1. Dirichlet multinomial mixtures using probabilistic modelling in R were used to stratify patients based on their microbial communities (operational taxonomic unit (OTU)-level) (23). The Laplace approximation was used to find the model of best fit and determine the number of clusters from the dataset. Unique microbial states were labelled Dirichlet clusters (DC). None of the measured clinical factors were significantly different between the two DC groups (DC1 and DC2) and disease controls (Table S2). A similar analysis was performed on phenotypic subtypes of CRS (CRSsNP, CRSwNP, CRSwCF) and disease controls (Table S3), where age and polyposis were found to be significant factors.

### Overall bacterial community composition

Quality filtering resulted in 346,003 bacterial 16S rRNA gene sequences from 31 samples. Samples that did not meet a rarefaction threshold of 677 were removed from further analysis, including 3 CRSsNP samples. A final number of samples included in further analyses was thus 8 disease controls and 20 CRS patients (CRSsNP = 5, CRSwNP = 8, CRSwCF = 7; DC1 = 13, DC2 = 7). The final, rarefied dataset included 200 taxonomically assigned OTUs at 97% sequence similarity (ranging from 4-45 OTUs per sample) and this OTU table was used for all subsequent microbial-related analyses.

Large inter-personal variation in microbial community composition was observed between individuals of each cohort (Fig. 1). The bacterial taxa *Staphylococcus* (OTU2), *Streptococcus* (OTU7), *Propionibacterium* (OTU6) and *Corynebacteriaceae* (OTU10) were prevalent in majority of the samples but only at low relative sequence abundances (Fig. 2A).

**Fig. 1:**
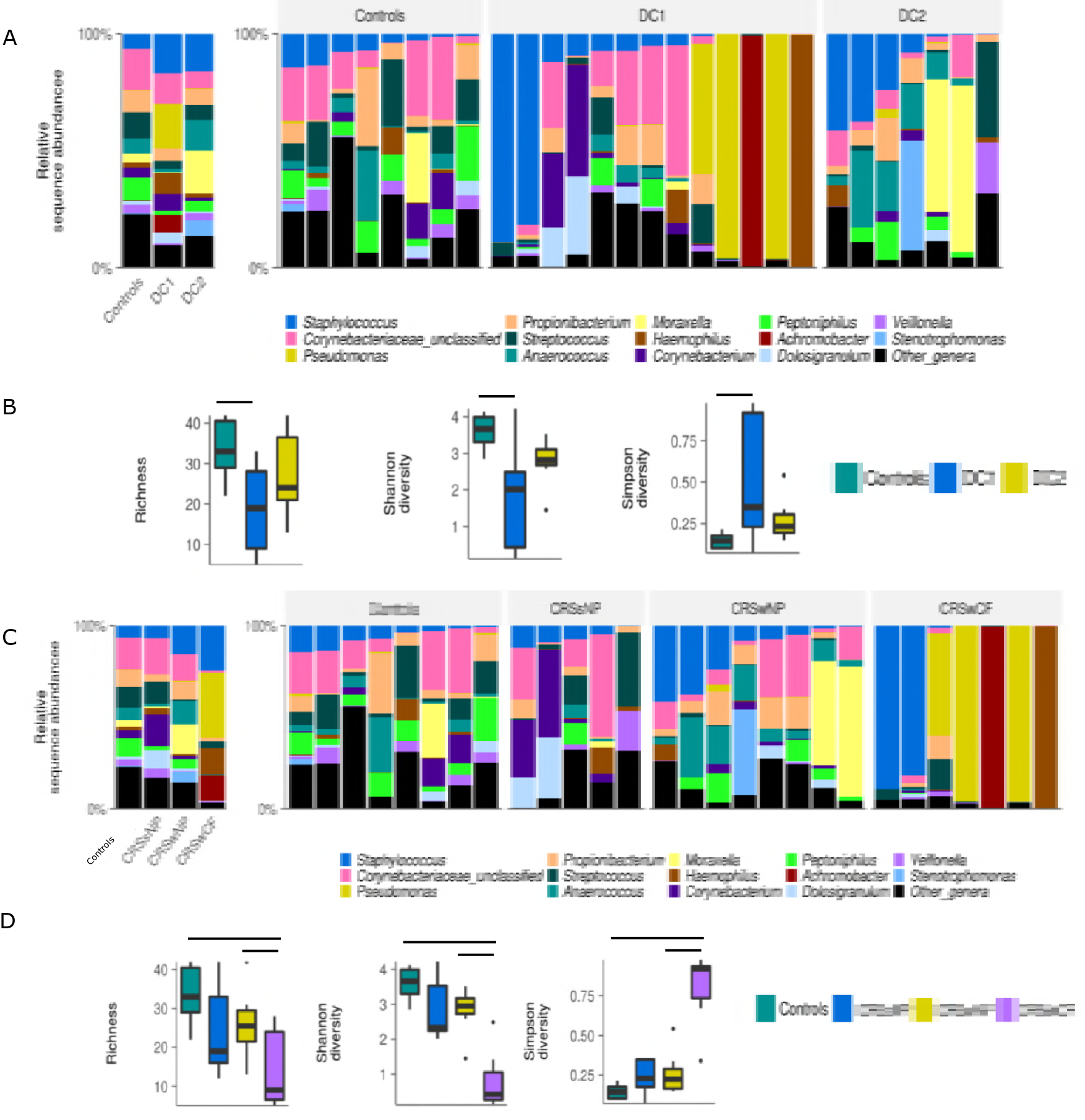
Bacterial community composition and alpha diversity for the CRS cohorts and disease controls. Grouped and patient-level profiles at genus-level are shown in the bar graphs for (A) DC groupings and (C) phenotypic grouping. Box-and-whisker plots represent group summaries for bacterial richness, Shannon diversity and Simpson diversity for (B) DC groupings and (D) phenotypic groupings. Horizontal lines represent significant differences (*p* < 0.05) between cohorts.

**Figure 2:**
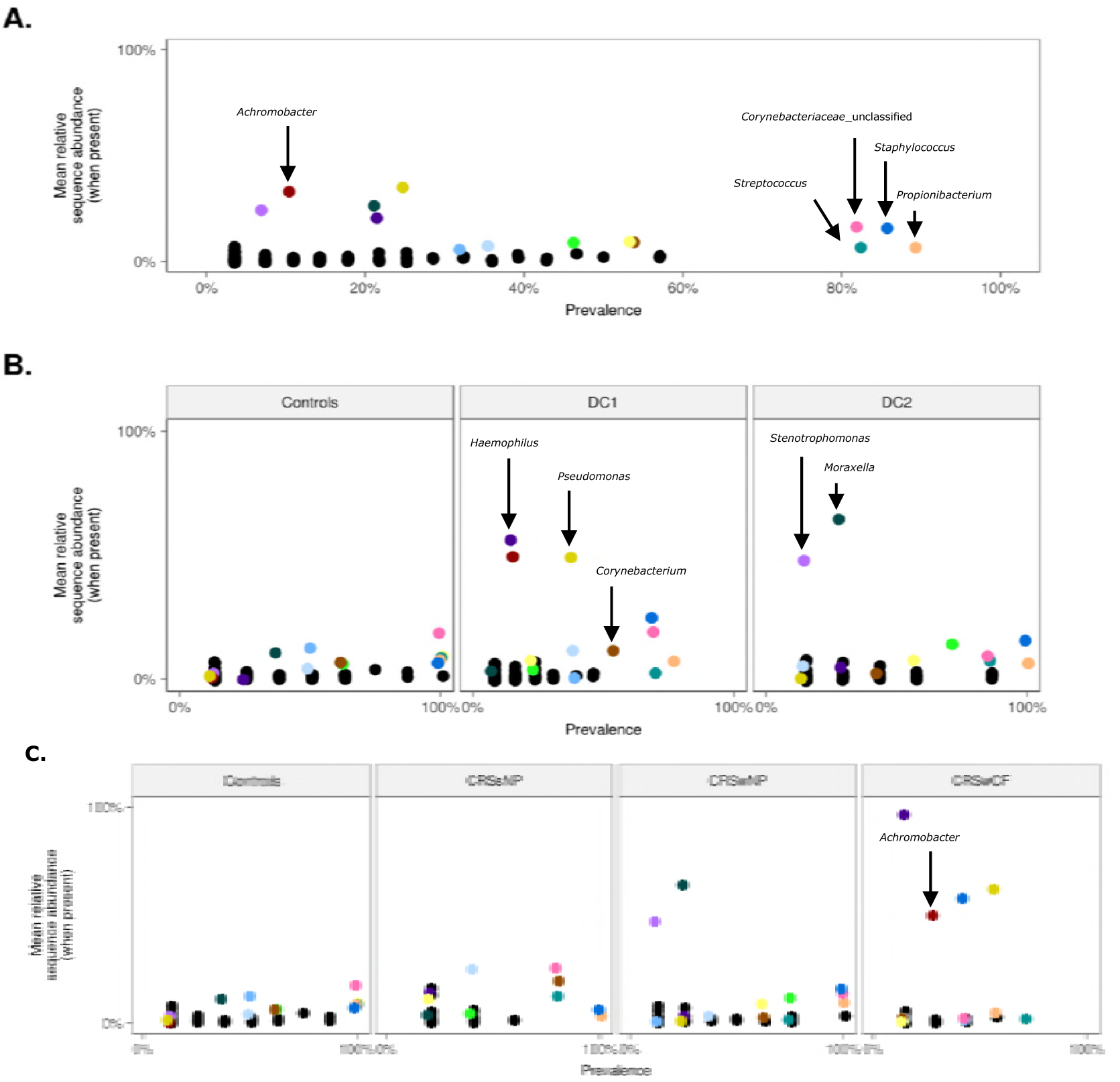
Mean OTU relative abundance per sample (when present in a sample) plotted against prevalence (occurrence) across (A) all samples, (B) each DC and control, and (C) each phenotypic CRS subtype and control. All OTUs are plotted. The 14 most abundant OTUs are color-coded, and all other OTUs are presented as black dots.

### Bacterial community composition of DC groups

Samples were assigned into clusters based on bacterial community composition using Dirichlet distributions (23). Dirichlet cluster 1 (DC1) comprised samples from all three CRS sub-groups (CRSsNP = 4, CRSwNP = 2, CRSwCF = 7). In contrast, DC2 was dominated by CRSwNP samples (n = 6) along with one CRSsNP sample.

A variety of alpha-diversity measurements revealed significant differences between DC1 and controls, however no significant differences were noted between DC1-DC2 and DC2-controls (Fig. 1B). The bacterial taxa *Pseudomonas* (average relative abundance 19% ± SD 37%), *Achromobacter* (7.6% ± SD 27%), and *Haemophilus* (8.5% ± SD 27%) dominated DC1, while DC2 was dominated by *Moraxella* (18% ± SD 31.5%) and *Stenotrophomonas* (6.7% ± SD 17.7%), but these taxa all had low prevalence as shown in Fig. 2B. Interestingly, controls were not dominated by any single bacterial taxon but instead had a low abundance of multiple genera, some of which were highly prevalent.

Taxa that were significantly different between groups were investigated through multiple pairwise comparisons at OTU- and genus-level (Table S4). OTUs that were significantly elevated in DC2 compared with DC1 were *Anaerococcus* (OTUs 12, 19 and 251), *Prevotella* (OTU200), *Megasphaera* (OTU179), and *Atopobium* (OTU203). Control samples were differentiated from DC1 and DC2 by significant increases in *Streptococcus* (OTUs 7, 18), *Veillonella* (OTU34), *Massilia* (OTU77), *Peptoniphilus* (OTU11) and *Halomonadaceae* (OTU109).

The dispersion of samples based on microbial community profiles in each DC was compared by calculating the distances to the centroid in non-metric multidimensional scaling (nMDS) analyses. DC1 samples were significantly more dispersed than control samples (*p* = 0.011, Tukey’s honest test) (Fig. 3B). In addition, Permutational Multivariate Analysis of Variance (PERMANOVA) test confirmed that each of the clusters identified for the CRS cohort and control group explained a significant proportion (R^2^ = 12.2%, *p* = 0.014) of the overall variation in the bacterial dataset. However these PERMANOVA results should be interpreted with caution in light of the different beta-dispersion patterns of the groups assessed (24).

**Fig. 3:**
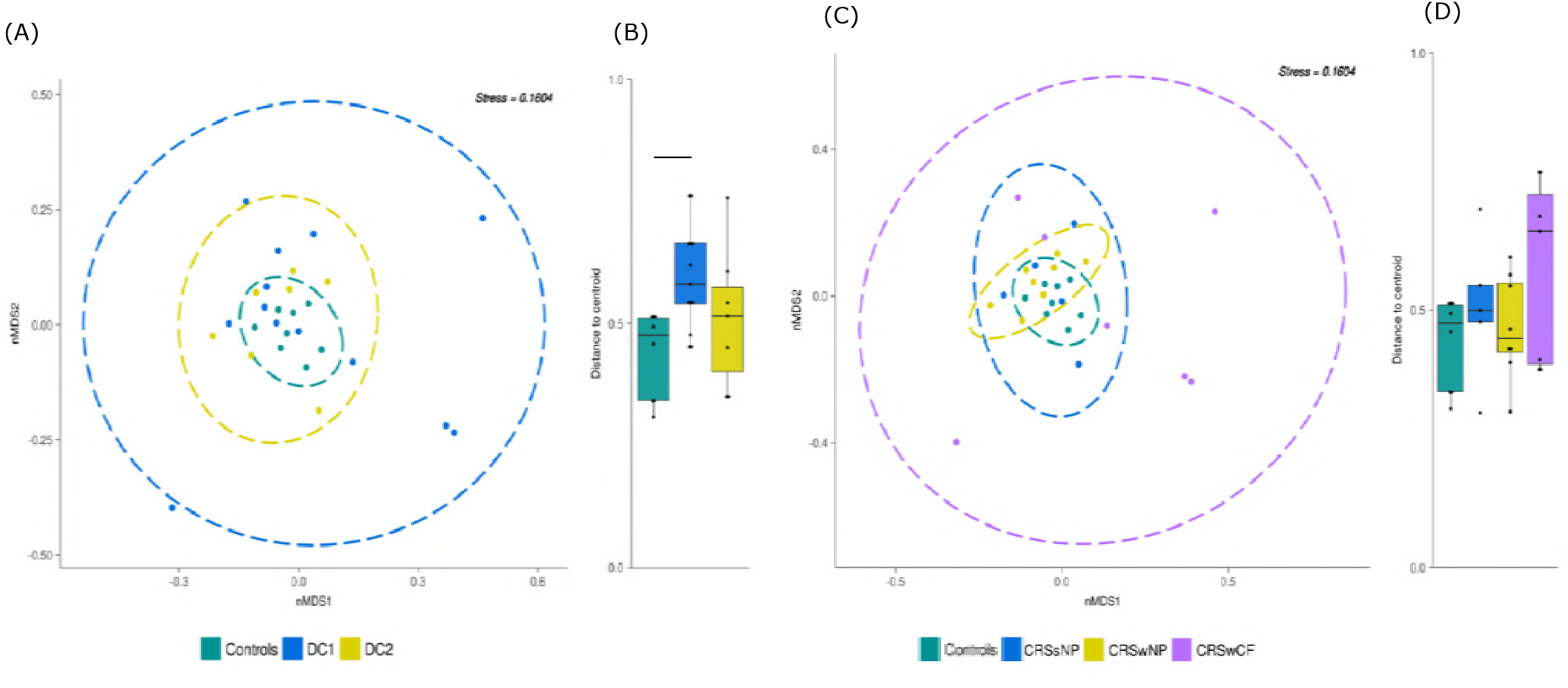
Non-metric multidimensional scaling plot using Bray-Curtis dissimilarity distances (weighted) for all samples for (A) DC clustering and (C) phenotypic subtyping. Ellipses represent the 95% CI spread from centroids. Box-and-whisker plot of distances between each subject to the centroid of their respective group (B) and (D). Beta-dispersion was significantly different between DC1 and controls (*p* = 0.0114; Tukey’s multiple comparisons of means).

### Bacterial community composition of phenotypic groups

Patients were also stratified based on clinical presentation of the disease, which includes the presence or absence of nasal polyposis and comorbidity of cystic fibrosis.

Alpha diversity was significantly (*p* <0.001) lower in CRSwCF patients compared with controls. Significant differences in diversity were also observed (*p* <0.05) between CRSwCF and CRSwNP samples.

Similar to controls (as described above), CRSsNP did not show dominance of any single bacterial taxon. However CRSwNP samples were dominated by *Moraxella* (average relative abundance 16% ± SD 30%) and *Stenotrophomonas* (5.8% ± 16.5%), while CRSwCF samples were dominated by *Pseudomonas* (35.4% ± 46%), *Staphylococcus* (24.5% ± 41.6%), *Achromobacter* (14.1% ± 37%), and *Haemophilus* (13.9% ± 36.8%) (Fig. 2C). Pairwise comparisons between control samples and CRS subtypes (CRSsNP, CRSwNP, CRSwCF) were performed on OTU- and genus-level data. CRSwCF samples were significantly reduced in *Propionibacterium*, *Corynebacterium*, *Anaerococcus* and *Peptoniphilus* compared with controls. Furthermore, CRSsNP samples had significantly lower abundance of *Peptoniphilus*, while CRSwNP were reduced in *Streptococcus* and *Veillonella*, compared to controls. OTUs and genera that were significantly different between CRS subtypes can be found in Table S5.

There were no significant differences in dispersion between phenotypic groups and controls when calculating analysis of variance. However, PERMANOVA tests were able to explain a larger proportion of the variation (R^2^ = 23.4%, *p* = 0.001) by phenotypic subtyping methods for CRS than with DC clustering.

### Tight junction protein and gene expression patterns in sino-nasal tissue

Based on our initial aim to investigate the underlying drivers of each subtype in CRS, the expression of 42 tight junction genes in sino-nasal tissue biopsies was measured for each patient and compared to that of disease controls. After results were normalised to a housekeeping gene, fold-changes in gene expression were recorded for each DC and CRS phenotypic subtype compared to controls.

Ten tight junction genes were significantly under- or over-expressed in DC1 and/or DC2 compared to controls (Fig. 4A). The gene ACTA1, which encodes a skeletal α-actin protein that maintains the cytoskeleton and cell movement, was the only gene to be significantly over-expressed in both DC1 and DC2. Each DC group exhibited unique tight junction gene expression patterns. Three genes were significantly under-expressed in DC1: CSDA, TCF7 and PVRL1. In DC2, nine genes were significantly under-expressed compared with controls (Fig. 4A).

**Figure 4:**
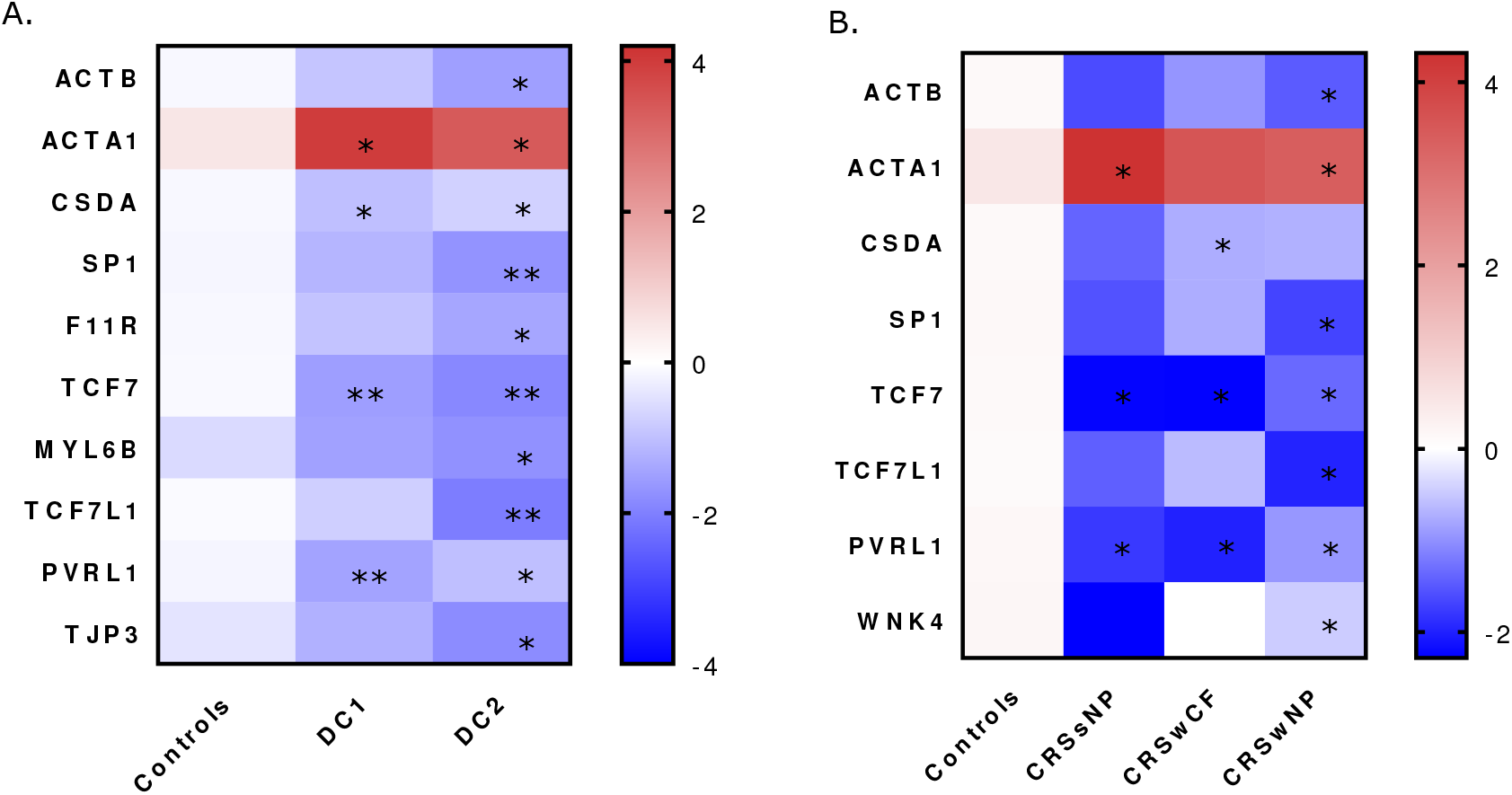
Tight junction genes that were significantly different in expression for (A) DC groups and (B) phenotypic subgroups compared with controls are displayed in the heatmap. Blue represents genes that have low level expression, and red represents higher expression. Values displayed are log-transformed mean fold-change (2^-ΔΔCt^). & represents *p* values below 0.05 and && represents *p* values below 0.01.

Subtyping patients based on phenotypic characteristics found 7 tight junction genes to be under expressed and ACTA1 gene to be over expressed in the CRS cohorts compared with the controls (Fig 4B). Genes PVRL1 and TCF7 were significantly under-expressed in all three CRS subtypes. The expression profile of CRSsNP and CRSwCF were very alike, with the only one exception (gene CSDA) which was under-expressed in CRSwCF patients (Fig. 4B). Spearman correlation analysis of the dataset after adjustment for multiple comparisons showed a significant negative association between tight junction gene MTDH expression and members of the bacterial genus *Pseudomonas* (Fig. S1).

Staining of sino-nasal tissue biopsies identified ZO-1 and occludin proteins at apico-lateral contact points of adjacent epithelial cells (Fig. S2). Claudin-1, by contrast, was predominantly seen in subapical regions, with concentrated, continuous staining amongst the mid-basal region of epithelial cells (Fig. S2). ZO-1 area staining was significantly (*p* = 0.03) lower in DC2 (mean 0.5 ± S.D 0.3%) compared with controls (1.1 ± 0.5%) (Table S3). There were no significant differences in claudin-1 and occludin staining in disease controls relative to DC1 and DC2. In addition, there were no significant differences in the staining of ZO-1, claudin-1 and occludin proteins in the three CRS subtypes compared with controls.

Furthermore, no correlations were observed between tight junction proteins and abundance of bacterial taxa (Fig. S1).

### Inflammatory state and mucosal integrity of tissue biopsies

Inflammatory marker cells (T-cells, B-cells and macrophages) were enumerated in the tissue biopsies to assess the inflammatory state of the patients. Although all three cell types were reduced in the disease controls compared with DC1 and DC2 cohorts, B-cells (CD20+) were the only identified inflammatory marker with a significant difference between the groups, with higher numbers in DC2 (Table S2).

In contrast, by traditional CRS subtyping approaches, all three inflammatory marker cells were significantly elevated only in CRSwNP cohort compared with controls. There were no significant differences between the CRS sub-type cohorts, except for the elevated amounts of macrophages (CD68+) in CRSwNP compared with CRSwCF.

The sino-nasal tissue biopsies were also assessed for mucosal integrity. Goblet cell enumeration and cilia integrity in the biopsy specimens were significantly reduced in DC2 compared with DC1 and control (Table S2). Similarly, when subtyping the same group of patients based on phenotypic approaches, goblet cell enumeration and cilia integrity were significantly reduced in CRSwNP patients compared with controls and CRSsNP patients. Interestingly, CRSwCF patients had no significant differences in mucosal integrity (collagen content, goblet cell enumeration and cilia integrity) compared with controls. Furthermore, Spearman correlation analysis showed a positive association between goblet cell counts and tight junction gene LEF1 expression (Fig. S1). No other significant associations were found between mucosal integrity, inflammatory marker cells and all other measured variables in this study.

## Discussion

Endotyping of CRS patients has been the subject of considerable recent research, as it is hoped that a sub-classification of this condition will allow for more specific and effective therapies to be administered. In this study, we defined microbial states for CRS using probabilistic modelling, in which patients with similar microbial states were clustered together. Furthermore, the same cohort of patients were also sub-typed based on phenotypic presentation of the disease. We then sought to understand the underlying influences on these cohorts by investigating sino-nasal mucosal integrity, tight junction gene/protein expression and inflammatory status. The two approaches of clustering CRS patients will be compared and discussed further.

## Resolving the microbial heterogeneity of CRS

Phenotyping of CRS patients based on clinical factors can be subjective and provides little information about microbes and their involvement in this disease. As shown previously for gastrointestinal, lower and upper respiratory diseases (11, 16, 17, 23), distinct microbial states were identified for CRS patients, allowing for stratification based on bacterial composition. The advantage of the new clustering approach used in this study and by others (11) is that it reflects a patient’s microbial state at the time and places the patient into a distinct microbial cluster type. Appropriate targeted treatment strategies could then be prescribed for patients in the future based on their distinct microbial pattern.

In this study, two distinct microbial states of CRS patients were identified that were significantly different to each other in diversity, beta-dispersion and the relative abundance of members from the genus *Anaerococcus*. Although this novel way of classifying CRS samples could explain some of the microbial variations in the dataset, there was still a large proportion of variation that could not be accounted for. This is most likely due to the sizeable inter-patient variation observed in this study and reported previously (15, 25). We anticipate with larger cohort sizes these microbial states would be more pronounced and less obscured by individual variation. While studies that use less invasive approaches to obtain samples such as nasopharyngeal swabs and rinses are able to increase cohort sizes (26, 27), these samples are not suitable for analyses of histology and mucosal integrity. In addition, tissue biopsies are required to study host gene expression levels. For these reasons, tissue biopsies were used in the analyses within this study.

Subtyping patients based on the phenotype of the disease (with or without nasal polyps or with cystic fibrosis) has been the most common approach to date. The results of the microbial communities from each of these subtypes is confirmatory to those previously described by our group (28, 29). Interestingly, a greater amount of the variation (23.4%) observed in the dataset from this study could be explained through this patient phenotypic clustering approach rather than by the DC approach.

## Potential drivers of microbial signatures for CRS patients

Understanding the underlying factors that shape the sino-nasal microbial community of a CRS patient is essential for defining the pathogenesis of this complex disease and for developing better treatment protocols. The most common hypothesis for the pathogenesis of CRS is that the epithelial barrier is defective (30). Allergens (such as pollen), host genetics, bacteria, viruses or inflamed tissue could all contribute to the disruption of the sino-nasal epithelial barrier (31). In this study, we chose to investigate several of these potential host factors that could contribute to CRS pathogenesis and determine their influence on the sino-nasal microbial state. Out of the 42 measured tight junction genes, nine exhibited reduced expression in CRS patients compared with controls. Reduced expression of tight junction genes in CRS patients compared with controls have been observed previously (6). The reason for the overexpression of ACTA1, which encodes for proteins involved in cell motility, structure and integrity (www.uniprot.org) is unclear but presumably reflects the host’s response to an impaired epithelial surface. Each DC in this study had a unique tight junction gene expression profile, suggesting that the two identified CRS DCs are functionally different from each other. It remains unknown whether specific tight junction gene expression could play a role in determining distinct microbial states in the nasal cavity, or vice versa. The significant association between tight junction genes and bacterial community composition observed in this study needs further validation with *in vitro* tests. However, this observation does provide some evidence of interactions between host tight junction gene expression and sino-nasal microbial communities.

Stratification of patients based on the traditional phenotypic approach were not able to clearly separate out the tight junction gene expression profiles of CRSsNP and CRSwCF cohorts. This lack of clarity, would suggest that future studies studying tight junction gene expression profiles in CRS patients should consider using alternative patient stratification approaches.

Of the three measured tight junction proteins in this study, only ZO-1 was significantly downregulated in DC2 relative to controls. ZO-1’s adhesive function between transmembrane proteins and the underlying actin skeleton denotes its critical role at the epicentre of the tight junction complex (32, 33). Furthermore, depletion of ZO proteins in mammary epithelial cells results in failure of tight junction strands to assemble and consequent loss of barrier function (33). These findings suggest an important role for ZO-1 in the establishment of functioning tight junction complexes. It is possible that reduced expression of ZO-1 protein could underlie subsequent downregulation of integral tight junction proteins, resulting in a loss of barrier properties. A loss of claudin-4, ZO-1, occludin and E-cadherin in tissue biopsies of CRS patients has been found previously (6, 34). The lack of any significant observations on the tight junction protein measurements in the phenotypic CRS sub groups compared with controls, also emphasises the need to look at other ways to stratify CRS. Another potential driver of microbial states in CRS is mucosal integrity. Several studies have reported significant changes in the structure and composition of the mucosa in CRS (35-37). CRS subjects exhibited polarising results in regards to evidence of mucus hypersecretion, with a significantly higher goblet cell count compared with controls. Interestingly, DC2 cohorts had significantly lower goblet cell counts and reduced integrity of cilia compared with DC1 and controls. This observation was also made for subgroup CRSwNP patients compared with other CRS cohorts and controls. Mucociliary function in CRS is essential for the physiological function and immunity of the nose (38). A loss in function promotes formation of biofilms and bacterial infections. Certain species of *Pseudomonas, Haemophilus* and *Streptococcus* produce ciliostatic or ciliotoxic agents that could result in loss of ciliary function (39). In this study, correlations with cilia integrity and goblet cell numbers and bacterial taxa were not observed. However, further research into toxin production by signature members of each cohort that results in cilia loss is required.

Previous studies have shown pro-inflammatory cytokines (IFN-γ and IL-4) to disrupt epithelial integrity *in vitro*, which in turn could cause changes to the sino-nasal micro-environment by disrupting surfaces for attachment or increase the permeability of microbes to the underlying tissue (6, 40). In addition, Cope et al. (11) found distinct microbial clusters to have unique patterns of immune response. Evidence from these studies suggests that microbial states are influenced by various host factors, possibly including cellular junctions. Furthermore, consistent with findings from other groups, CRS patients in this study based on microbial states (DC1 and DC2) or phenotypic subtyping (CRSwNP) had higher abundance of B-cells in the mucosa, compared with controls. Previous efforts have shown a proliferation of B-cells in the sino-nasal mucosa when exposed to antigens (41). Inflammatory cytokine IL-13 is a key factor in stimulating antibody production (such as IgE and IgA) by B-cells in response to antigen exposure in airway inflammatory diseases (42). IgE levels are elevated in eosinophilic CRSwNP patients (43). Accordingly, the elevated levels of B-cells in CRS patients is a good indication of the inflammatory status of the sino-nasal tissue of CRS patients.

## Stratification approach for CRS

The traditional approach of stratifying CRS patients based on their phenotypic presentation of the disease is standard practice, however such approaches are struggling to unravel the complexities of the disease. This study showed that, while phenotypic subtyping approaches can help explain some of the variation in the microbial community dataset, alternative methods using microbial signatures can also have some benefit. Each microbial state of CRS in this study was found to have a unique tight junction expression profile, which was not observed for the phenotypic subtyping approach. Accordingly, it remains unclear whether microbial states influence, or are being influenced by, barrier impairment. To answer these questions, *in vitro* studies will need to be undertaken in the future.

## Materials and Methods

### Patient recruitment and sample collection

Twenty-three adult patients undergoing functional endoscopic sinus surgery for CRS by a single surgeon (RD) were recruited for this study. Diagnosis and subsequent recruitment of CRS patients were based on EPOS 2012 guidelines (44). These included CRS patients with polyps (CRSwNP; n = 8), CRS patients without polyps (CRSsNP; n = 8), and CRS patients with cystic fibrosis (CRSwCF; n = 7). Control subjects (n = 8) without any signs of sinus disease undergoing endoscopic surgery for the removal of pituitary tumours or medial orbital decompression were also recruited. Control patients exhibited no evidence of mucosal inflammation on endoscopy or computed tomography scans. Exclusion criteria included patients being administered systemic corticosteroids or antibiotics within four weeks prior to surgery, age less than 18 years, immunodeficiency, pregnancy, and other comorbidities (apart from cystic fibrosis). This study was approved by the Health and Disability Ethics Committee of New Zealand (NTX/08/12/126). Prior to sample collection, informed written consent was obtained from all patients.

Patients completed a symptom score sheet, prior to surgery, in which the following CRS symptoms were rated on a scale of 0-5: nasal obstruction, anterior nasal discharge, posterior nasal discharge, facial pain or fullness, and loss of smell. These scores were summated to give the ‘Symptom Severity’ score in Table S1. Lund-Mackay scoring was used to quantify radiological disease severity (45). Tissue biopsies were collected intraoperatively from the ethmoidal sinus under general anaesthesia prior to administration of topical vasoconstrictors or intravenous antibiotics. Biopsied specimens were rinsed in sterile saline and partitioned. Samples for quantitative PCR and bacterial community analysis were fixed in RNAlater for 24 h, then stored at −20°C. Samples for immunohistochemistry analysis were fixed in Carnoy’s solution (60% ethanol, 30% chloroform, 10% glacial acetic acid) before paraffin embedding.

### Bacterial community analysis

#### DNA extraction

DNA was extracted from tissue biopsies using sterile Lysing Matrix E bead tubes (MP Biomedicals, Australia) and the AllPrep DNA/RNA Isolation kit (Qiagen, Germany) as previously described (46). A negative DNA extraction control using 200 μL sterile water was carried out simultaneously.

#### Bacterial community sequencing

The V3-V4 region of the bacterial 16S rRNA gene was amplified using primers 341F and 806R (47), with Nextera DNA library Prep Kit adapters attached. PCR reactions, amplification conditions and purifications were carried out as previously described (12). In brief, genomic DNA (~100 ng) from each sample was amplified in duplicate PCR reactions of 35 cycles then pooled to a final volume of 50 μL. Negative PCR controls were included in all PCR reactions and yielded no detectable amplicons. Eluent from the negative extraction control was also subjected to PCR amplification and yielded no detectable product. Purification using Agencourt AMPure magnetic beads (Beckman Coulter Inc., USA) was carried out as per the manufacturer’s instructions. Purified PCR products were quantified using Qubit dsDNA High-Sensitivity kits (Life Technologies, New Zealand), standardised to ~5 ng per sample, and submitted to the University of Auckland Genomics Centre for library preparation and sequencing using Illumina MiSeq (2 x 300 bp paired-end reads). Raw sequence reads were deposited into the SRA-NCBI database (BioProject ID: PRJNA482256).

#### Bioinformatics

Sequences were merged and quality filtered in USEARCH (version 8.0) with default settings as previously described (12). OTU clustering based on a 97% 16S rRNA gene sequence similarity threshold was performed using the UCLUST algorithm in USEARCH (48). Each OTU was taxonomically assigned in QIIME (49) using the RDP classifier 2.2 against the SILVA 16S rRNA gene database (version 128) (50, 51). Sequences mapping to the human genome were removed from subsequent analyses. Samples were rarefied to an even sequencing depth of 677 reads. Alpha-diversity (including Shannon, Simpson, and richness (observed OTUs)) and beta-diversity (including weighted and unweighted UniFrac distances, and Bray-Curtis dissimilarity) were calculated in QIIME. Graphical outputs were created in R (version 3.4.1) (52) and GraphPad Prism (version 7.03).

#### RNA extraction and quantitative PCR (qPCR)

RNA extraction of biopsied tissue was carried out in parallel to the DNA extraction using the AllPrep DNA/RNA Isolation Kit (Qiagen) as per the manufacturer’s instructions. Extracted RNA was transferred to a collection tube and re-eluted in 30 μL DNA/RNA-free sterile water.

Recovered RNA was treated with DNaseI (Invitrogen) for selective degradation of contaminant DNA as per the manufacturer’s instructions. The quantity and quality of RNA were measured using a Nanodrop 3000 spectrophotometer. Successful removal of genomic DNA from DNase-treated RNA samples was demonstrated by PCR targeting the human beta-actin gene (53). DNase-treated RNA (standardised to ~100 ng/μL) was converted into cDNA using iScriptTM Reverse Transcription Supermix for RT-qPCR (Bio-Rad, New Zealand) as per manufacturer’s instructions.

Pre-designed, 384-well qPCR arrays (TJ H384) were obtained from Bio-Rad Laboratories Inc. (Auckland, New Zealand) for analysis of 42 known tight junction genes and one house-keeping gene (GAPDH). Each sample was assessed for PCR performance, reverse transcription efficiency, DNA contamination, and RNA quality. Analysis of results was carried out using the ABI Prism 7900HT detection system (version 2.4). Each sample was run in duplicate on separate qPCR arrays and averaged for further analysis. Tight junction gene expression was normalised to the house-keeping gene GAPDH. Mean fold-change of gene expression in CRS-affected patients compared to control was calculated using the equation 2^-ΔΔCt^ (54).

### Histological analysis of sino-nasal tissue

Paraffin-embedded tissues were prepared into 4 μm-thick sections then mounted on Superfrost Plus positively-charged microscope slides (Thermo Fisher Scientific). Detailed procedures for staining and analysis of tight junctions (ZO-1, claudin-1 and occludin), goblet cells, cilia, collagen and inflammatory cells (CD4: T cells, CD20: B cells, CD68: macrophages) can be found in the Supplementary Material.

### Statistical analysis

Statistical analyses were carried out using GraphPad Prism software (CA, USA) and R. Dirichlet multinomial mixtures using probabilistic modelling in R was used to stratify patients based on their microbial communities (OTU-level) (23). The Laplace approximation was used to find the model of best fit and determine the number of clusters from the dataset. Unique microbial states were labelled Dirichlet clusters (DC). Sample dispersion between groups was compared using analysis of variance, permutation test (PERMDISP), and Tukey’s honest significant differences in R. The software package ‘adonis’ PERMANOVA using distance matrices (Bray-Curtis dissimilarity (weighted and unweighted), UniFrac distances (weighted and unweighted)) was used to compare between centroids of groups.

Multiple non-parametric pairwise comparisons for categorical variables (diagnosis, gender, ethnicity, polyposis, smoking status, antibiotic and steroid usage, and co-morbidities) were tested using Fisher’s exact test, with Bonferroni adjustment for multiple comparisons. Pairwise comparisons between continuous variables for bacterial diversity, age, Lund-Mackay scores, symptom severity scores, tight junction gene expressions, histological analyses, bacterial OTUs and genera were tested using Dunn’s test, with Bonferroni adjustment for multiple comparisons. Heat maps depicting Spearman correlations, with significance calculated using Spearman coefficients, including multiple comparisons adjustment using Benjamini & Hochberg False Discovery Rate (BH-FDR) with hierarchical clustering of correlation coefficients were generated in R. Factors with significant (*p* < 0.05) positive or negative correlations were plotted.

## Acknowledgments

The authors would like to thank the Garnett Passe and Rodney Williams Memorial Foundation for its generous support of this research. Authors declare no conflict of interest. Authors K.B., M.W.T, R.G.D contributed to the study design. Authors K.B., R.C., S.G., M.H, S.W-T., J.H., K.C., B.WM contributed to data collection and analysis. Authors K.B., R.C., M.W.T., R.G.D. contributed to manuscript preparation.

